# Retinoid-X Receptor Agonists Increase Thyroid Hormone Competence in Lower Jaw Remodeling of Pre-Metamorphic *Xenopus laevis* tadpoles

**DOI:** 10.1101/2021.11.30.470596

**Authors:** Brenda J. Mengeling, Lara F. Vetter, J. David Furlow

## Abstract

Thyroid hormone (TH) signaling plays critical roles during vertebrate development, including regulation of skeletal and cartilage growth. TH acts through its receptors (TRs), nuclear hormone receptors (NRs) that heterodimerize with Retinoid-X receptors (RXRs), to regulate gene expression. A defining difference between NR signaling during development compared to in adult tissues, is competence, the ability of the organism to respond to an endocrine signal. Amphibian metamorphosis, especially in *Xenopus laevis*, the African clawed frog, is a well-established in vivo model for studying the mechanisms of TH action during development. Previously, we’ve used one-week post-fertilization *X. laevis* tadpoles, which are only partially competent to TH, to show that in the tail, which is naturally refractive to exogenous T3 at this stage, RXR agonists increase TH competence, and that RXR antagonism inhibits the TH response. Here, we focused on the jaw that undergoes dramatic TH-mediated remodeling during metamorphosis in order to support new feeding and breathing styles. We used a battery of approaches in one-week-old tadpoles, including quantitative morphology, differential gene expression and whole mount cell proliferation assays, to show that both pharmacologic (bexarotene) and environmental (tributyltin) RXR agonists potentiated TH-induced responses but were inactive in the absence of TH; and the RXR antagonist UVI 3003 inhibited TH action. At this young age, the lower jaw has not developed to the point that T3-induced changes produce an adult-like jaw morphology, and we found that increasing TH competence with RXR agonists did not give us a more natural-metamorphic phenotype, even though Bex and TBT significantly potentiated cellular proliferation and the TH induction of *runx2*, a transcription factor critical for developing cartilage and bone. Prominent targets of RXR-mediated TH potentiation were members of the matrix metalloprotease family, suggesting that RXR potentiation may emphasize pathways responsible for rapid changes during development.

## Introduction

An organism’s acquired ability to respond both qualitatively and quantitatively to a physiological signal, defined as competence, is distinguished between endocrine signaling during development, which tends to lead to irreversible, organizational effects, from that of healthy adult tissues, which controls the functioning of tissues and organs to maintain homeostasis. Thyroid hormone (TH) action regulates many aspects of vertebrate development including cartilage growth and skeletogenesis (1–4). Over developmental time, the vertebrate organism traverses from low to high TH competence (5). Vertebrate development depends upon appropriate timing and concentrations of TH for good biological outcomes. During human development, adverse outcomes arise from both insufficient and excessive TH (1,6–10). However, analysis of the effects of TH on mammalian development are confounded by maternal effects due to the nature of intrauterine growth. Amphibian metamorphosis, the process through which larval tadpoles develop into adult frogs, overcomes these obstacles; it provides an accessible and dramatic model for direct investigation of the role TH plays during vertebrate development (11–14). Metamorphosis is initiated and maintained through the action of TH (15–18). The African clawed frog, *Xenopus laevis*, provides an accessible and effective laboratory model for assessing the role of TH throughout development, and its metamorphosis has been shown to model the essential perinatal surge in TH signaling in humans (11,14).

In all vertebrates, TH acts through the thyroid hormone receptors (TRs), which are DNA-binding, ligand-regulated transcription factors of the nuclear receptor (NR) superfamily (19,20). THs are identical across all taxa, and the TRs are highly conserved between *X. laevis* and humans (12,13). Two isoforms of TR are expressed from two different genes, TRα and TRβ. In *X. laevis* tadpoles, TRα is expressed before synthesis of THs commences (21), whereas TRβ expression is induced after the nascent thyroid gland begins to synthesize THs through TH binding to TRα; it is a direct target gene of TRs (22,23). 3,3’,5-triiodo-L-thyronine (T3) is the TH with the highest affinity for the TRs (24–26). TRs bind DNA and can regulate gene transcription in both the absence and presence of TH. In the most-studied model of TH action, apo-TRs recruit co-repressor proteins, which close the chromatin environment to the general transcription machinery, causing transcriptional repression. In contrast, T3-bound TRs recruit co-activator proteins, which open the chromatin environment to the general transcription machinery, causing transcriptional activation (19,20). TRs heterodimerize with another NR, the retinoid-X receptors (RXRs) (27). RXRs bind several natural ligands, including 9-cis retinoic acid, and they can dimerize with many different NRs in addition to TRs (28). The TR-RXR heterodimer shows higher affinity for DNA, especially in the presence of T3, than the TR-TR homodimer (29).

Due to the importance of TH signaling for proper development, man-made chemicals that disrupt TR action have the potential to produce adverse outcomes (30). Our initial investigations into disruption of TR function used an integrated TRE-driven Luc reporter in a rat pituitary cell line that endogenously expresses both TRα and TRβ (31,32). In that cell culture system, vitamin A (VA) metabolites were the primary hits for disrupting the TRE-Luc reporter (33). VA metabolites can activate both the retinoic acid receptors (RARs) and the RXRs, depending upon the metabolite (34). RARs, like TRs and RXRs, are members of the NR superfamily. The documented but not mechanistically understood ability of VA metabolites to convert into each other make using them to determine RAR vs RXR function difficult. A pharmaceutical RAR ligand did not affect the TRE-Luc reporter (32), but a pharmaceutical RXR agonist did (33). Since RARs should not bind a TRE, and RXRs heterodimerize with TRs on TREs, these results would appear unsurprising were it not for the fact that, in most adult tissues and cells, RXR ligands are unable to affect the action of the TR-RXR heterodimer (35,36). Pituitary cells, like the reporter cell line, are an exception, wherein RXR ligands do affect the ability of TR to control the hypothalamus-pituitary-thyroid (HPT) axis (37); the biological reasons for this are not understood. In fact, the pharmaceutical RXR agonist used in this study, bexarotene (brand name Targretin, Bex), produces severe hypothyroidism in patients given the drug, which limits its usability as a chemotherapeutic (38–40). However, given the inability of RXR ligands to affect TR function in peripheral tissues, such as the liver—a major site of TR function—the TR-RXR heterodimer is generally considered to be an example of a “non-permissive” RXR heterodimer, meaning that only the ligand for the TR, TH, can induce activation. This is also thought to “make sense” because TH is an endocrine hormone that tightly regulates several important biological functions like heart rate, and tight, endocrine-type regulation would be complicated if a second hormone or environmental ligand (dietary, etc) were also able to affect TR-RXR action (35).

Tributyltin (TBT) is a pervasive environmental pollutant that was the first described endocrine disruptor when it was discovered that exposure to TBT caused marine gastropods to develop imposex phenotypes, where female gastropods develop male secondary sex characteristics (41,42). TBT was widely used as an antifoulant in marine paints. Mechanistic work determined that TBT functioned through the mollusk RXR, and that treating marine gastropods with either 9-cis retinoic acid or TBT produced the same imposex phenotype (43–45). Biophysical investigations showed that TBT covalently binds to a cysteine residue at the entrance to the RXR ligand-binding pocket, creating an activated conformation in the RXR (46). In the rat pituitary reporter cell line, TBT behaved like Bex, strongly suggesting that it was functioning as an RXR agonist (33,47). These results left open the question as to whether RXR agonists in a developing organism would behave like RXR agonists in most adult tissues (i.e. RXR agonists have no effect on TR action) or like in our pituitary reporter cell line (i.e. RXR agonists could modulate TR function).

Previously, we developed a suite of quantitative assays to assess function and possible disruption of TH action in 1-week post-fertilization (1wk-PF) tadpoles (NF 48) (48). 1wk-PF tadpoles express TRα, but they do not yet have an active thyroid gland; therefore, they are TH negative and are considered pre-competent (21). Addition of T3 to their rearing water activates many metamorphic pathways, but the addition of T3 does not make their TH competence complete. For example, tail resorption, the last step of metamorphosis, is minimal even under supraphysiological doses of T3, because at least in part, the tail expresses high levels of the T3-deactivating enzyme, deiodinase 3 (49). We found that the addition of Bex or TBT to the rearing water of 1wk-PF tadpoles in the presence of T3 significantly potentiated the action of T3 in larval tissues undergoing resorption, including the tail (47,50). In effect TBT/Bex increased T3 competence in the tail to near metamorphic levels. At the transcriptomic level in the tadpole tail we found that TBT acted identically to Bex, solidifying that the mechanism of TBT action on TH function was at the level of RXR agonism (50). We wondered if the potentiating effects of RXR agonists affected metamorphic phenotypes beyond resorption.

Amphibian metamorphosis affects almost every tissue system and cell fate decision. TH induces the jaw to remodel to facilitate the transition from an herbivorous tadpole to a carnivorous adult frog. During natural metamorphosis, visible jaw morphological changes start at NF 59, which is approximately 45 days post-fertilization (PF) under ideal rearing conditions (51,52). Thomson describes three phases of Meckels cartilage development in the lower jaw (LJ): 1) a lag phase (NF 57-59) with low levels of cell proliferation, 2) a division phase (NF 60-62) of rapid cell division, and 3) a synthesis phase (NF 62-66) wherein the matrix content of the cartilage increases significantly (53,54). Rose showed that tadpoles prior to NF 57 (∼41 days PF) respond to the TH but the beak-like morphological changes that result are not seen in a natural metamorphosis (51). Between NF 48 and NF 57 significant, non-TH-induced growth occurs to the cartilages of the lower jaw, and this growth appears to be essential for producing appropriate morphology upon TH administration. Bearing this in mind, we investigated whether RXR ligands were able to potentiate the T3-induced changes that are possible at NF 48, where we have an extant suite of quantitative assays to monitor potential disruption of T3 action (48). We found that both Bex and TBT potentiated T3-induced proliferation, the activation of *runx2*, a transcription factor necessary for maturation of cartilage and bone ossification, and the matrix metalloproteases *mmp11* and *mmp13l*. On the other hand, the RXR antagonist UVI 3003 (UVI) (55) prevented T3-induced morphological changes while not inhibiting proliferation, and it only selectively inhibited gene transcription. In addition, Bex and TBT still potentiated T3 action in the LJ in tadpoles at NF 54, which are considered prometamorphic and fully competent to respond to THs.

## Materials and Methods

### Reagents

3,3’,5-triiodo-L-thyronine (T3, T6397-100MG) and tributyltin chloride (TBT, T50202-5G) were purchased from MilleporeSigma (Burlington, MA) and Bexarotene (Bex, 5819/10 and UVI 3003 (UVI, 3303/10) were purchased from Tocris Biosciences (Bio-Techne, Minneapolis, MN). All treatment ligands were dissolved or diluted in dimethyl sulfoxide (DMSO, Thermo Fisher Scientific, Waltham, MA). oLH (ovine luteinizing hormone) was purchased through the National Hormone and Peptide Program (Los Angeles, CA), pregnant mare serum gonadotropin was purchased from Thermo Fisher Scientific, and tricaine methanesulfonate was purchased from Western Medical Supply (Arcadia, CA).

### Animal husbandry

The laboratory has an approved University of California Davis Institutional Animal Care and Use protocol that covers the husbandry and mating of adult *Xenopus laevis* frogs and ligand exposure of larval tadpoles. Wild-type *X. laevis* frogs were mated and embryos cultured as described (Mengeling 2017).

### Tadpole precocious metamorphosis morphology assay

NF 48 (1-week post-fertilization) tadpoles were treated, fixed for photography, and dorsal head photos taken using a Leica DFC3000 G camera on a Leica MZLFIII microscope as described (Mengeling 2016 and 2017). Treatment concentrations, unless otherwise indicated were 10 nM T3, 30 nM Bex, 1 nM TBT, and 600 nM UVI, based upon previous results. The angle of the lower jaw was measured using the FIJI (56) distribution of ImageJ (57). GraphPad Prism 9 (GraphPad Software, La Jolla, CA) was used to generate box and whisker plots, where boxes represent the 25^th^ to 75^th^ percentiles with the bar at the median, and whiskers are maximum and minimum values. For statistical analyses, each animal counted as an individual, and 2 clutches (ten tadpoles/clutch) were assayed independently to control for clutch-to-clutch variability. NF 54 tadpoles were treated as NF 48 animals except that the volume/tadpole of rearing water was increased to 50 ml, and treatments were stopped at three days rather than five, due to the extreme gill resorption in the T3 + Bex animals. Three independent clutches of NF 54 tadpoles were used with 4-5 tadpoles per clutch.

### Immunohistochemistry of lower jaws for proliferation

The lower jaws from tadpoles fixed as for morphology were removed as follows: a straight cut was made just posterior to the olfactory epithelium and anterior to the eyes. the upper and lower jaw were separated, and two diagonal cuts were made on the outer rim of the jaw to separate the cartilage from the excess tissue. LJs were treated as described for immunohistochemical analysis of phospho-Histone H3 reactivity (EMD Millepore, 06-570, 1/300 dilution) (48,58). Anti-phospho-Histone H3 (Ser10) was from EMD Millepore (06-570, 1/300 dilution), and goat anti-rabbit IgG (H+L) conjugated with Alexa Fluor 488 was from Molecular Probes (A11008, 1/400 dilution). Positive cells were counted from blinded images using the Cell Counter tool of Fiji.

### Gene Expression

Tadpoles were treated with ligands for 48 hours as for morphology and as described (47,50), using a 2-way ANOVA design: vehicle (DMSO), T3, RXR ligand, and T3 + RXR ligand. Lower jaws were isolated from unfixed tadpoles as for immunohistochemistry. Pools of 15 LJs from a single clutch were used for total RNA extraction. LJ tissue was disrupted and homogenized by bead beating with two 0.125-inch stainless steel beads for 1 minute in a Mini-Beadbeater-16 (Biospec Products, Bartlesville, OK). Total RNA was extracted using the RNeasy Plus Mini Kit per the manufacturer’s instructions (Qiagen, Germantown, MD). Total RNA was quantified using a NanoDrop (Thermo Fisher Scientific, Waltham, MA). One microgram of total RNA was used to synthesize cDNA with the High-Capacity Reverse Transcription Kit (Thermo Fisher Scientific), and 0.5 μl of cDNA from a 20-μl reaction was used in a 10-μl reaction using PowerUp SYBR Green Master Mix (Thermo Fisher Scientific) in a Roche LightCycler 480. The *X. laevis rpl8* gene was used as a normalizer. Statistics were performed using 2-way ANOVA analysis with a Sidak’s multiple comparison test (MCT) in GraphPad Prism 9. Sequences for the primers used for quantitative PCR are given S1 Table.

### Transgenic tadpole luciferase reporter assay

NF 54 tadpoles, sorted at 1wk-PF for GFP+ expression in the eye lens, were staged by assessing morphology of the hind limb, according to the normal scale by Nieuwkoop and Faber (59), and then treated through their rearing water for two days as previously described (50). Treatment concentrations were 10 nM T3 and 2 nM TBT. No mortality arose from the treatments over the treatment period. After treatment, tadpoles were anesthetized in 0.1% MS-222 (Western Medical) buffered with 0.1% sodium bicarbonate. The LJs were excised and minced on ice prior to freezing and then processed and assayed as described. Each animal was treated as an individual for statistical purposes (n = 9 per treatment) from two independent clutches (4 animals in one clutch and 5 in the other). Two-way ANOVA using clutch and treatment as covariates with Tukeys MCT to compare treatments was used from GraphPad Prism 9.

## Results

### RXR agonists potentiate T3-induced morphological changes to the lower jaw, and an RXR antagonist abrogates T3 effects

Using our precocious metamorphosis assay system, we treated *X. laevis* 1wk-PF tadpoles (NF 48) for five days by exposure through their rearing water with vehicle or 10 nM T3 in the presence or absence of RXR ligands. This treatment period did not result in animal mortality under any of the treatment conditions. Previously, we found that 30 nM Bex and 1 nM TBT produced maximal, non-toxic responses, and so we used them here (47,50). The dorsal head photos in Figure 1 show representative animals from each treatment regimen. Vehicle treatment resulted in normal tadpole morphology (Figure 1a), and treatment with the RXR ligands in the absence of T3 (Figure 1b-1d) did not result in morphological changes. Treatment with 10 nM T3 (Figure 1e) resulted in visible gill resorption and decreased the angle of Meckels cartilage. Co-treatment with either RXR agonist, Bex or TBT, potentiated the T3-inductions of gill resorption and the angle of Meckels cartilage (Figure 1f-1g). However, co-treatment with the RXR antagonist UVI 3003, abrogated the effect of T3 on both morphologic phenotypes.

**Figure 1:**
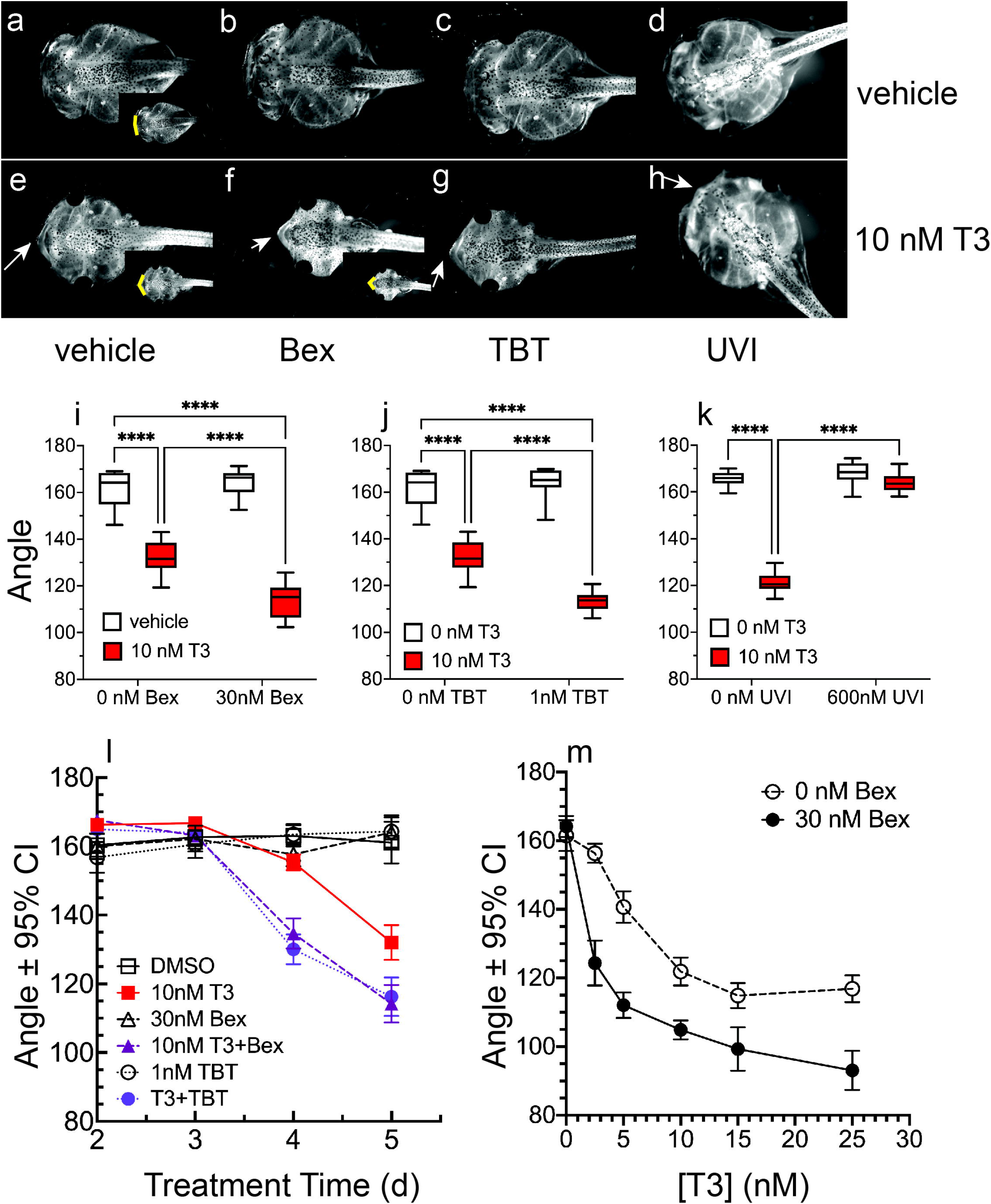
RXR agonists potentiate T3-induced changes to lower jaw morphology, while an RXR antagonist abrogates T3 action. a-h: Representative dorsal head photos of tadpoles treated for five days starting at 1wk-PF (30 nM Bex, 1 nM TBT, 600 nM UVI). i-k: Quantification of changes to the jaw angle. Boxes represent 25^th^-75^th^ percentiles with the line at the median (n = 10-15 from 2-3 clutches), and whiskers represent the min and max values. Statistics show results from Sidak’s multiple comparison test in conjunction with 2-way ANOVA (****, p < 0.0001). l: Effect of Bex and TBT on T3-induced jaw angle changes as a function of time. Data points represent means from 20 animals from two different clutches; error bars delineate the 95% confidence intervals, indicating statistical significance. m: Treatment with 30 nM Bex augments jaw angle narrowing as a function of T3 dose. Statistics are the same as in the time course, although the clutches were different.

In order to quantify the effects of T3 and the RXR ligands on Meckels cartilage, we measured the angle of the LJ (Figure 1i-k) from independent clutches of tadpoles, using ten animals per clutch. The inset photos (Figure 1a, 1e, 1f) show the change in angle that was measured. Protrusion of the Meckels cartilage caused a decrease in the LJ angle. Figure 1i shows that in the presence of T3, 30 nM Bex significantly potentiated the decrease in the LJ angle (compare red boxes). 2-way ANOVA analysis indicated significance for the interaction between T3 and Bex (p < 0.0001. As with our study on the effects of RXR agonists on T3-induced tail resorption, 1 nM TBT behaved almost identically to 30 nM Bex; the interaction between T3 and TBT was significant (p < 0.0001). In contrast, co-treatment of T3 and the RXR antagonist UVI prevented T3 action, and the LJ angle was not significantly changed from vehicle-treated tadpoles (Figure 1k); however, due to the strong abrogation of the T3-induction by UVI, the interaction between T3 and T3+UVI was still significant by 2-way ANOVA (p < 0.0001). Figure 1l shows the LJ angle measurement as a function of treatment time. Again, co-treatment of either Bex or TBT with T3 caused an identical response that showed an acceleration of the Meckels cartilage protrusion. Tadpoles treated for four days with T3 plus RXR agonist had the same decrease in LJ angle as tadpoles treated for five days with T3-alone. Over a T3-dose curve, the T3-induced decrease in LJ angle was significant starting at 5 nM T3 (error bars represent the 95% confidence interval), and all doses of T3 in the presence of Bex showed a significantly reduced LJ angle compared to T3-alone, such that 5 nM T3 plus Bex/TBT produced the same LJ angle as 15 nM T3 (Figure 1m), which is the dose that produces the maximal change in LJ angle.

### RXR agonists potentiated T3-induced cellular proliferation in Meckels cartilage, but the RXR antagonist had no effect

In young tadpoles, exogenous T3 administration triggers proliferation in several tissues, including the LJ (58). We excised LJs after four days of treatment for whole mount immunohistochemistry (IHC) of the mitotic marker phosopho-Ser10 Histone 3 (pH3) to assess the effects of T3 and RXR ligands on cellular proliferation in Meckels cartilage. 1wk-PF tadpoles were treated for four days instead of five to facilitate LJ removal; T3-induced changes to the gills and brain make removing the LJ more difficult after five days of treatment. Proliferative cells were counted from blinded images over the area of Meckels cartilage (Figure 2a). Figure 2b-f show representative photos of different ligand treatment combinations from which proliferative cells were counted. For quantification, each combination of T3 and RXR ligand were assayed with two independent clutches, and for each clutch, RXR agonist potentiation was significant. Figure 2g-i shows the two clutches combined for each group. Vehicle-treated LJs had few proliferative cells (Figure 2g-i). In contrast, treatment with 10 nM T3 increased the number of mitotic cells at least 15-fold for each treatment group. Co-treatment of either 30 nM Bex (Figure 2g) or 1 nM TBT (Figure 2h) RXR agonists with the T3 resulted in a significant increase in the number of proliferating cells in the Meckels cartilage. Since the RXR agonists induced a significant increase in proliferative cells, we expected that co-treatment of the RXR antagonist UVI with T3 would result in a decrease in proliferative cells. However, as Figure 2i shows, that is not the case; UVI did not inhibit cellular proliferation in Meckels cartilage. Therefore, the RXR agonists potentiated both the decrease in LJ angle and the induction of cell division; however, UVI prevented T3 action morphologically (Figure 1h) but had no effect on T3-induction of proliferation (Figure 2i). Aurora kinase B (*aurkb*) is the kinase that performs the phosphorylation of Ser10 of H3. T3 induced *aurkb* mRNA expression (figure 2j-l); however, neither Bex (Figure 2j) nor TBT (Figure 2k) significantly increased that induction, suggesting that increased *aurkb* expression alone was not the mechanism through which the RXR agonists potentiated cell proliferation in Meckels cartilage. UVI inhibition of *aurkb* was not significantly different from T3-alone (p = 0.081) (Figure 2l).

**Figure 2:**
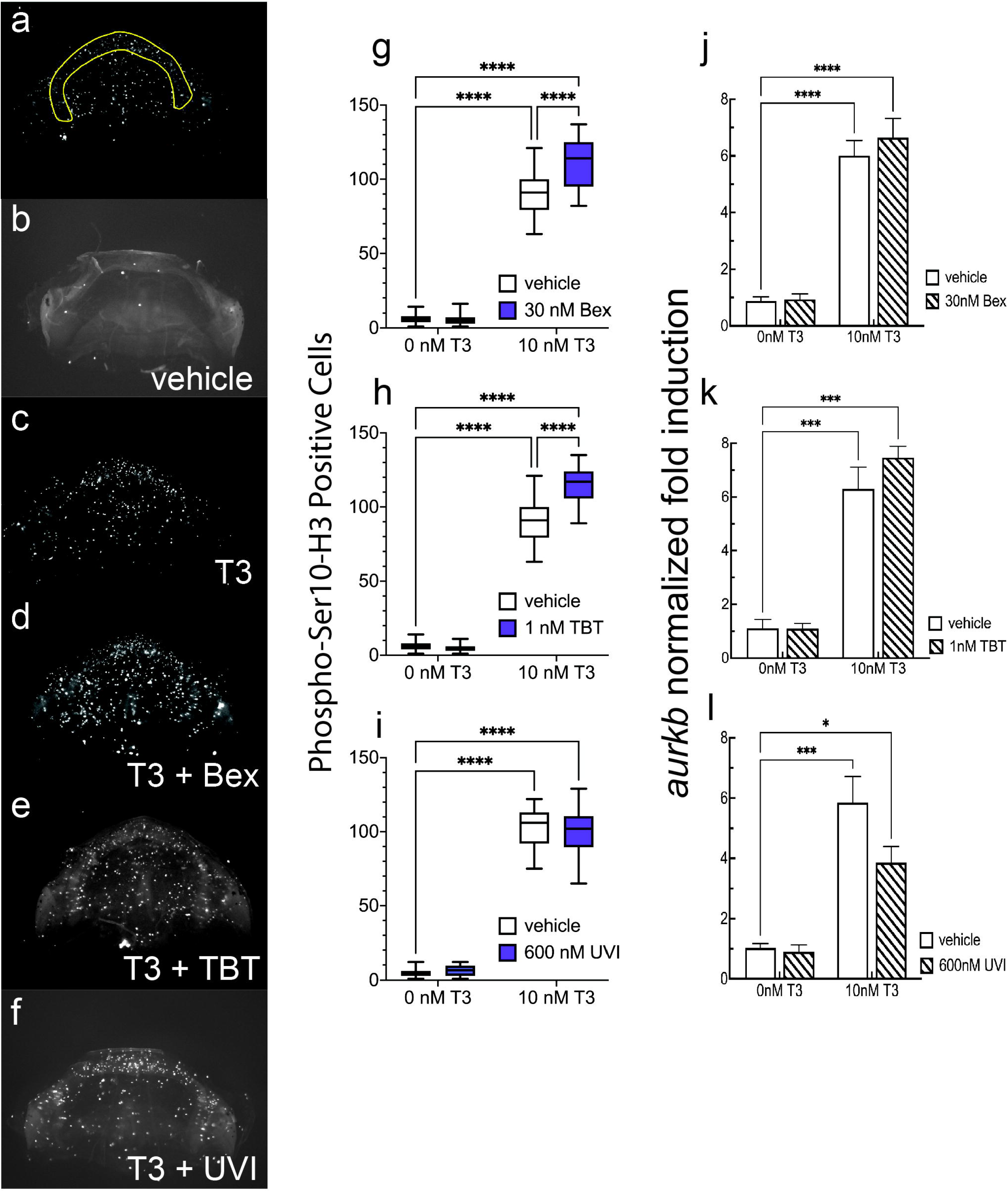
RXR agonists potentiate T3 action on cellular proliferation in the LJ of 1wk-PF tadpoles. a: Region of Meckels cartilage used for quantitation of proliferation. b-f: Representative photos of the effects of different treatments on proliferation using phopho-Ser10-H3 reactivity. g-i: Quantification of proliferation in the presence and absence of T3 and RXR ligands. Boxes and statistics are as in Figure 1 (n = 20-30 jaws from 2-3 clutches). j-l: RXR ligands do not significantly affect the T3-induced expression of aurora kinase B mRNA (*aurkb*). Bars represent the mean of 3-6 independent clutches, and statistics show results from Sidak’s multiple comparison test in conjunction with 2-way ANOVA (****, p < 0.0001; ***, p < 0.001; *, p < 0.05).

### RXR agonist potentiation of gene expression is gene specific

Our previous work examining the role of RXR ligands to perturb T3-mediated gene expression in the tails of 1-wk-PF tadpoles after a 48-hour induction, showed that the *bona fide* TR target gene for TRβ, *thrb*, was modestly, but significantly, potentiated by the RXR agonists and inhibited by the antagonist when assayed at the transcriptomic level using Tag-Seq. However, over a time course assayed by RT-qPCR, the same two-day time point showed no significant potentiation and inhibition by the agonists and antagonist, respectively (50). Using RT-qPCR to assess *thrb* expression in the LJ after two days of treatment, we found significant activation by T3 (white bars in Figure 3a), but neither Bex nor TBT potentiated that induction (slashed bars in Figure 3a, Bex, TBT). UVI also did not inhibit the T3 induction (slashed bar in Figure 3a, UVI). TH-bZIP is a transcription factor that is one of the most strongly TH-induced genes during metamorphosis. It is encoded by the *thibz* gene, and it is another TR direct target gene, having at least two TREs in the promoter region (60). In the LJ, T3 strongly induced *thibz* expression (Figure 3b, white bars), but the RXR agonists did not potentiate the signal (Figure 3b, slashed bars, Bex, TBT). However, UVI did significantly reduce the T3 induction of *thibz* (Figure 3b, slashed bar, UVI). In the tail, we found the same outcome: the RXR agonists did not affect *thibz* expression, but the RXR antagonist significantly did (47,50). These results strongly suggest that the RXR agonists and antagonist are not always operating reciprocally.

**Figure 3:**
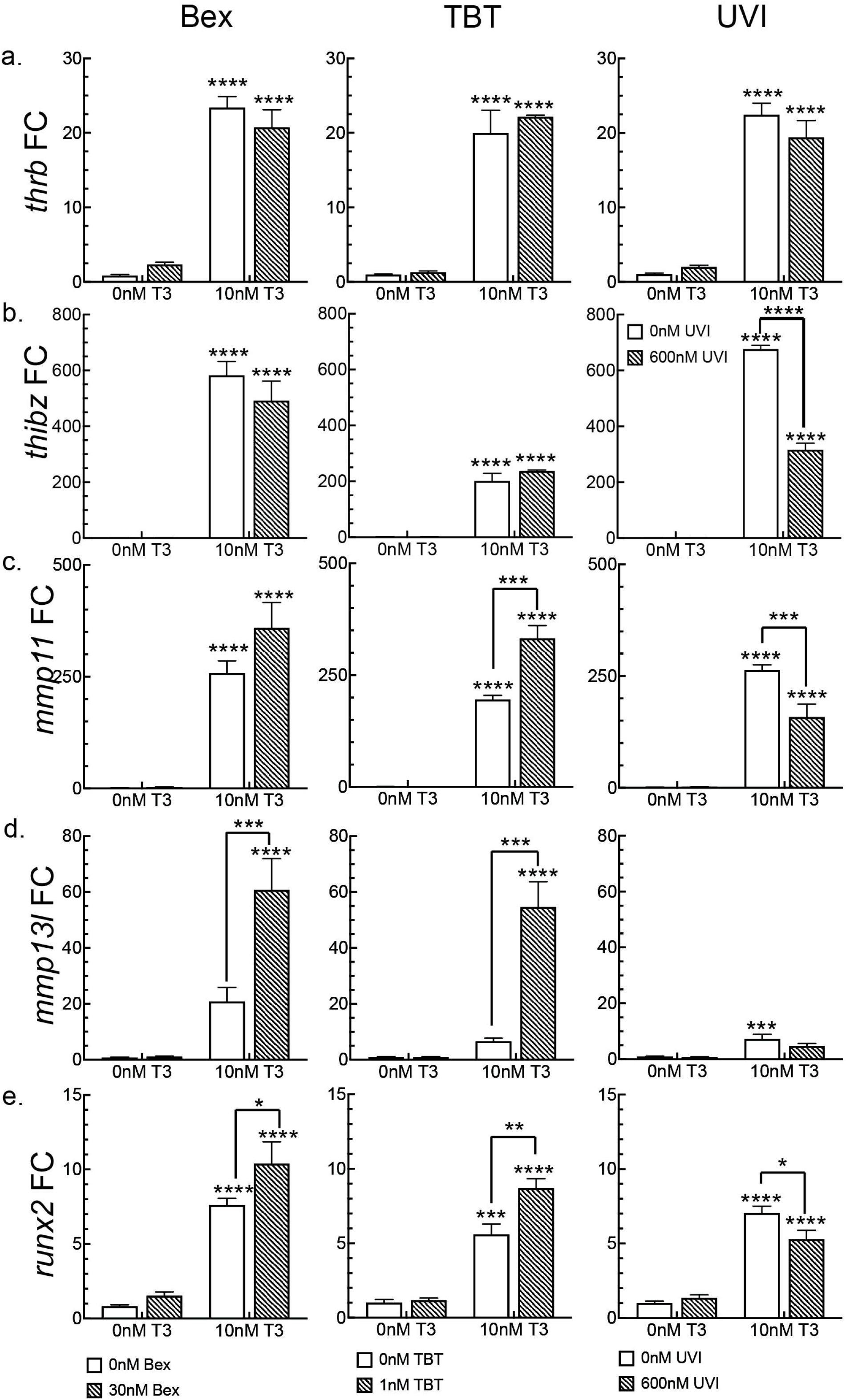
RXR ligands have gene-specific effects on T3-induced differential gene expression. Left column: The effect of RXR agonist Bex on T3-induced gene expression. Middle column: The effect of environmental RXR agonist TBT on T3-induced genes. Right column: The effect of RXR antagonist UVI on T3-induced genes. Striped bars indicate the presence of the RXR ligand, and white bars show induction in the absence of the RXR ligand. Statistics show results from Sidak’s multiple comparison test in conjunction with 2-way ANOVA (****, p < 0.0001; ***, p < 0.001; **, p < 0.01; *, p < 0.05).

During metamorphosis, matrix metalloprotease activity is essential for both tissue resorption and tissue remodeling. We and others have shown the importance of stromelysin-3 (*mmp11*) and collagenase-3 (*mmp13l*) expression (61–63). We found in the LJ that *mmp11* was strongly activated by T3 (white bars in Figure 3c). Co-treatment with 30 nM Bex increased *mmp11* expression, although this result did not reach statistical significance (p = 0.068), using the maximal number of biological replicates (n = 6) recommended for pooled, outbred animal tissues. However, co-treatment of T3 with 1 nM TBT did result in significantly potentiation of *mmp11* expression (n =3, p = 0.0004). Furthermore, UVI inhibited the T3 induction of *mmp11* significantly (n = 4, p = 0.001). In the experiments using Bex, even though T3 on its own activated the *mmp13l* gene 20.8-fold (S.E.M. = 4.95) (white bars in Figure 3d), this activation did not reach statistical significance (p = 0.0589), like it does in the tail. However, co-treatment with Bex increased *mmp13l* activation to significance (p < 0.0001) compared to both vehicle and T3-only treatments (slashed bar in Figure 3d, Bex). This situation held true for co-treating T3 with TBT (slashed bar in Figure 3d, TBT): T3-alone activation of *mmp13l* was not significant (p = 0.65), while T3 + TBT treatment was significantly potentiated (p < 0.0001) compared to both vehicle and T3-alone (slashed bar in Figure 3d, TBT). In contrast, in the experiments using UVI, the 7.3-fold activation by T3-alone did reach statistical significance (p = 0.0008, n = 4), but UVI did not significantly inhibit T3 induction of the gene (p = 0.14).

Runx2 is a transcription factor that is required for the transition from proliferating chondrocytes to hypertrophic chondrocytes in the maturation of cartilage for the development of a bony skeleton (64– 66). In non-amniote animals like fish and amphibia, it is required earlier for rostral cartilage formation (67,68). Due to the extensive changes to jaw cartilage during metamorphosis, we investigated whether T3 regulated its expression. In the LJ, T3 induced expression of *runx2* approximately 7-fold (white bars in Figure 3e), and this induction was significantly potentiated through co-treatment of either Bex or TBT with the T3 (slashed bars in Figure 3e, Bex, TBT). In addition, UVI co-treatment significantly inhibited *runx2* induction by T3 (slashed bar in Figure 3e, UVI). T3 did not regulate the expression of *runx3*, nor did we see activation of certain *runx2* downstream targets like *col10a1* (collagen10α1) (data not shown).

### RXR agonists potentiate T3-action in TH-competent (NF 54) tadpoles

While 1wk-PF tadpoles are considered only partially competent to respond to THs, tadpoles at NF 54 (approximately 26 days PF) are considered fully competent to respond to THs and to be entering metamorphosis (21,49,61). We raised tadpoles to NF 54, using hind limb development to determine the developmental stage (59), and then treated them with 10 nM T3 in the presence and absence of 30 nM Bex to investigate whether the RXR agonist could still potentiate the action of T3 in a fully competent tadpole. Tadpoles were treated for three days with compounds (a longer treatment time was not possible due to the extreme gill resorption in T3 plus Bex animals), and then we measured the LJ angle. Figure 4a (white boxes) shows that T3-alone caused a small but significant decrease in the lower jaw angle. As in NF 48 tadpoles, Bex-only treatment had no effect on the lower jaw morphology—tadpoles were indistinguishable from vehicle-treated. Bex co-treatment with T3 significantly potentiated the decrease in the LJ angle at this later stage of growth (Figure 4a), suggesting that the ability to increase the competence for T3 in the lower jaw was still possible, even for these presumed fully competent animals.

**Figure 4:**
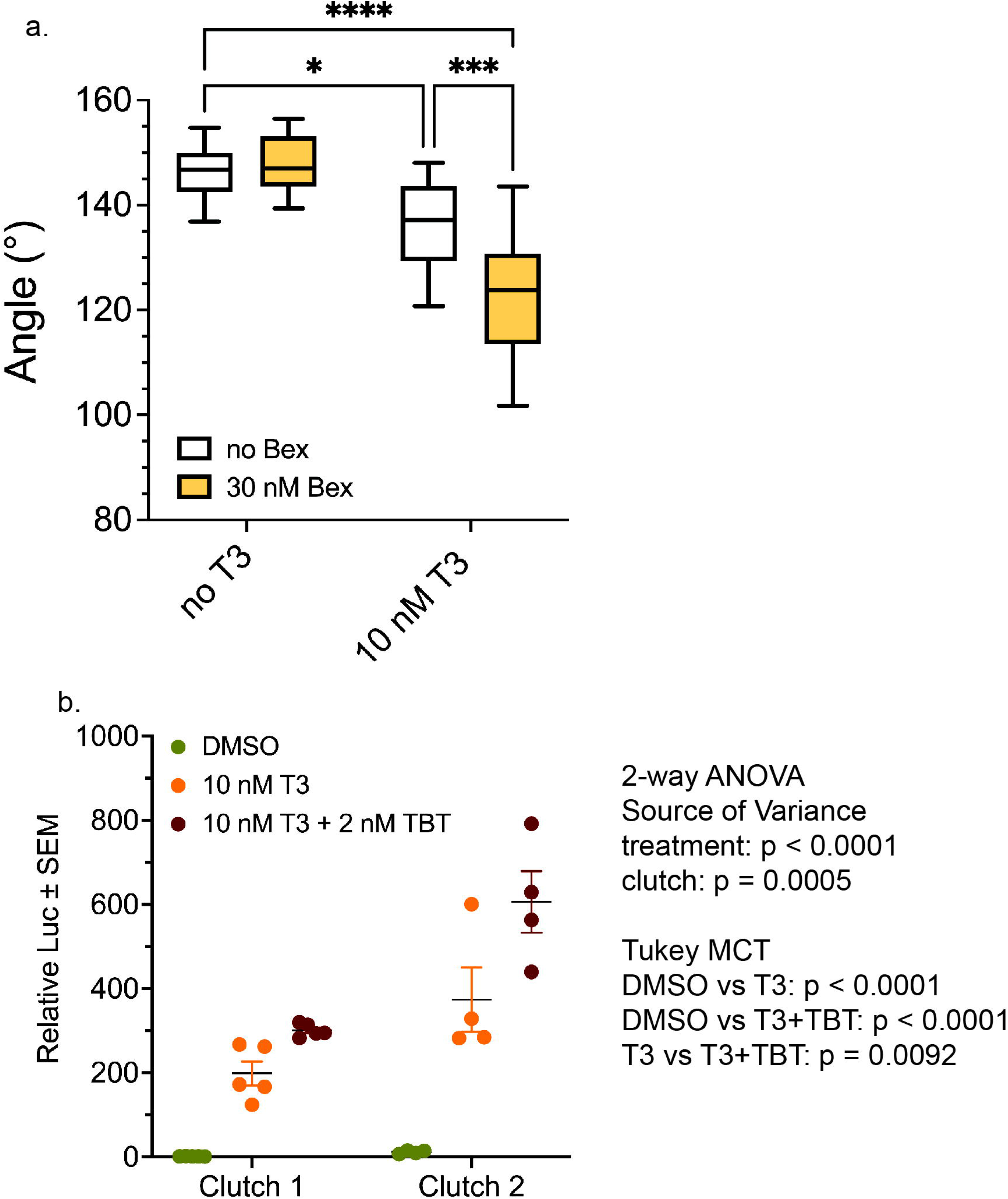
RXR agonists potentiate T3 action in the LJ in pro-metamorphic NF 54 tadpoles. a. Bex potentiates the T3-induced decrease in the LJ angle in NF 54 tadpoles treated for three days. Boxes and statistics are as in Figure 1 (n = 14 jaws from 3 clutches). Statistics show results from Sidak’s multiple comparison test in conjunction with 2-way ANOVA (****, p < 0.0001; ***, p < 0.001; *, p < 0.05). b. TBT potentiates T3-inducible, integrated luciferase reporter expression in the LJ of NF 54 tadpoles.

Previously we developed a transgenic line of *X. laevis* frogs that express firefly luciferase (Luc) under the regulation of the *X. laevis thibz* TH response elements (TREs) (47,48). In NF 48 tadpoles, assaying the entire head for Luc activation is required in order to generate a signal robust enough for statistics. At NF 54 we are able to analyze individual tissues, so we treated NF 54 tadpoles for 2 days with 10 nM T3 in the presence and absence of 2 nM TBT, and then we excised the lower jaws as we did for gene expression analysis. Luc activity was determined in the lower jaw samples and was normalized to the protein concentration of each sample. We assayed two clutches independently using two different TRE-Luc-bearing F2 male frogs to generate embryos with two different wild-type female frogs. TRE-Luc F2 males, even though they arise from the same founder female, display different levels of Luc activation by T3 that are nonetheless consistent within a clutch. Figure 4B shows the results of both clutches individually, showing the different levels of T3 activation between the two clutches. For clutch 2, a TBT-only treatment was also included and showed no activation. Using a 2-way ANOVA analysis of the combined data from both clutches where treatment and clutch were covariates, clutch was a significant source of variance (p = 0.0005), as was treatment (p < 0.0001). Using a Tukey multiple comparisons test post hoc on the combined clutch data, TBT significantly potentiated the T3 activation of the Luc reporter (p = 0.0092). This result indicates that the RXR agonists at this high-TH-competence stage could further increase the competence of LJ tissue for T3 at the beginning of natural metamorphosis.

## Discussion

In this report we have expanded upon our earlier findings concerning the ability of RXR agonists to function as a competence factor for TH signaling during vertebrate development (47,48,50). The poor biological outcomes that arise from insufficient or inappropriate TH during development have demonstrated the need for assessing the ability of man-made chemicals present in the environment to disrupt those signaling pathways.

In order to look at TH disruption in vivo and during development, we have used amphibian metamorphosis of the African clawed frog, *Xenopus laevis*. Metamorphosis performs two reciprocal functions: 1) development of adult tissues and organs required for life as a frog, and 2) removal of larval tissues no longer needed by the adult frog. Limb formation and growth and lung development are examples of development of new tissues and organs, and jaw development is an example of remodeling that must occur for the herbivorous tadpole to become a carnivorous frog. The other side of the metamorphic coin involves the resorption of larval tissues that are no longer required in the frog, such as gills and the tail. Naturally, removal of larval tissues must occur after the adult tissues have developed and become functional. For example, tail resorption is the last step in metamorphosis because it must occur after limb development is complete and the limbs are functional for locomotion. Under natural development, it takes approximately two months to go from a fertilized egg through a larval tadpole to a juvenile frog, with the metamorphic transition from tadpole to frog taking approximately 4.5 weeks under ideal conditions (13,18,21,59).

Our studies here employed a precocious metamorphosis assay, to determine whether a disruptor of TH signaling, which we have previously described disrupting larval tissue resorption phenotypes (47,50), can also disrupt a larval-to-adult remodeling function, namely, cartilage development in the LJ. By using 1wk-PF tadpoles, we were able to control the dose of TH, as tadpoles at this age do not yet synthesize THs. In this scenario, T3 and the potential disrupting chemicals taken up by the tadpole through administration in the rearing water. Although the LJ of the 1wk-PF tadpole is not able to support normal metamorphic changes to the LJ, molecularly the LJ can respond to T3 administration with reproducible morphological and molecular readouts.

Previously, we found that both the pharmaceutical RXR agonist Bex and the environmental RXR agonist TBT disrupted TH signaling in 1wk-PF tadpoles by significantly potentiating the ability of T3 to drive gill and tail resorption. Furthermore, the RXR antagonist UVI abrogated T3 action. Bex and TBT functioned identically in a global transcriptomic analysis of T3 signaling in the tail (50), indicating that TBT was functioning as a bona fide RXR agonist (43,44,46,69–72). Here, we show that the RXR agonists potentiate T3 action in the LJ by accelerating the rate of change and by increasing the potency of each T3 dose. As in the tail, TBT and Bex behaved nearly identically in the LJ independent of the experimental readout. In addition, the RXR antagonist abrogated the morphological changes induced by T3. We also measured the ability of the agonists and antagonist to disrupt T3-induced cellular proliferation. TBT and Bex both significantly potentiated proliferation, but UVI did not inhibit it. These findings suggest that the mechanisms of RXR agonist potentiation and of RXR antagonist inhibition are not strictly reciprocal. Furthermore, in contrast to their effects on proliferation, the opposite was seen in their effects on the T3-induction of the *thibz* gene. There, the RXR agonists had no effect, and the RXR antagonist significantly inhibited *thibz* activation. This was also seen in tail expression of *thibz* (50). How the agonists and antagonist are working at the molecular level is beyond the scope of these studies, but more than one mechanism is in play. Interestingly, RXR agonist s and the antagonist do not always behave in a reciprocal manner at all molecular or cellular targets when examined in detail.

UVI prevented morphological changes to the LJ in the presence of T3 but did not inhibit cellular proliferation, which suggests that cellular proliferation was not the main driver behind the morphological narrowing of the LJ. A better fit to the morphology patterns observed with RXR agonists and antagonist modulation of the LJ T3 response is the expression patterns of the matrix metalloproteases we tested. Both *mmp11* and *mmp13l* expression levels were potentiated by RXR agonists and inhibited by UVI. This was also true for the transcription factor *runx2*, which in mice is required for formation of ossified bones (73). In *Xenopus* and zebrafish, *runx2* is required earlier for cranial cartilage formation (67,68), but in our hands, it was significantly activated by T3 exposure, and that activation was potentiated by the RXR agonists and inhibited by UVI. We believe this is the first example of T3 activating *runx2* expression. In human thyroid cancer and breast cancer cells, TRβ suppressed the expression of *runx2* in the presence of TH, acting as a tumor suppressor (74,75).

An advantage of using 1wk-PF tadpoles for characterizing disruptors of TH signaling is the size uniformity of the tadpoles. We normally don’t have to normalize to the vehicle-treated control in each clutch, as we didn’t in Figure 1. However, as the tadpoles age, this size uniformity disappears, making morphological measurements more intrusive, as the animals must be housed separately and anesthetized and photographed before treatment for individual comparisons to after treatment changes. An advantage of assaying the LJ angle, is that it does not scale with tadpole head size; therefore, tadpoles can be group housed and measured only after fixation at the end of treatment. This provides a facile assay for TH disruption over developmental time, which in the case of RXR ligands, as they affect TH competence, could change as the animal develops and intrinsically increases in TH competence.

That said, we also chose NF 54 to assess whether the RXR agonists could still potentiate T3 action in the LJ because that is when plasma T3 is first detectable, and therefore, NF 54 is often considered the dividing line between premetamorphic and metamorphic tadpoles (21). However, NF 54 is nearly three weeks before metamorphic morphological changes in the jaw become apparent at NF 59 (51), and it is approximately two weeks before exogenous T3 leads to normal metamorphic development in the LJ. Therefore, TH competence in the LJ may still not be complete at NF 54 so that the cartilages can continue to develop in their normal T3-independent fashion until they are in the form that can remodel appropriately to an adult jaw. Thus, as prometamorphosis proceeds, the animal may be vulnerable to inappropriate RXR ligand activity from the environment. Ordinarily, endogenous retinoids can be controlled by the P450 retinoid-degrading enzymes, (76,77), yet organotins, or other as yet unknown chemicals in the environment that activate RXR, evade this buffer, and, therefore, still pose a unique and challenging problem for the exquisitely timed process of metamorphosis.

## Acknowledgements

The authors would like to acknowledge Michael L. Goodson for blinding the images for counting proliferative cells.

